# Enhanced selectivity of transcutaneous spinal cord stimulation by multielectrode configuration

**DOI:** 10.1101/2023.03.30.534835

**Authors:** Noah Bryson, Lorenzo Lombardi, Rachel Hawthorn, Jie Fei, Rodolfo Keesey, J.D. Peiffer, Ismael Seáñez

## Abstract

**Objective:** Transcutaneous spinal cord stimulation (tSCS) has been gaining momentum as a non-invasive rehabilitation approach to restore movement to paralyzed muscles after spinal cord injury (SCI). However, its low selectivity limits the types of movements that can be enabled and, thus, its potential applications in rehabilitation.

**Approach:** In this cross-over study design, we investigated whether muscle recruitment selectivity of individual muscles could be enhanced by multielectrode configurations of tSCS in 16 neurologically intact individuals. We hypothesized that due to the segmental innervation of lower limb muscles, we could identify muscle-specific optimal stimulation locations that would enable improved recruitment selectivity over conventional tSCS. We elicited leg muscle responses by delivering biphasic pulses of electrical stimulation to the lumbosacral enlargement using conventional and multielectrode tSCS.

**Results:** Analysis of recruitment curve responses confirmed that multielectrode configurations could improve the rostrocaudal and lateral selectivity of tSCS. To investigate whether motor responses elicited by spatially selective tSCS were mediated by posterior root-muscle reflexes, each stimulation event was a paired pulse with a conditioning-test interval of 33.3 ms. Muscle responses to the second stimulation pulse were significantly suppressed, a characteristic of post-activation depression suggesting that spatially selective tSCS recruits proprioceptive fibers that reflexively activate muscle-specific motor neurons in the spinal cord. Moreover, the combination of leg muscle recruitment probability and segmental innervation maps revealed a stereotypical spinal activation map in congruence with each electrode’s position.

**Significance:** Improvements in muscle recruitment selectivity could be essential for the effective translation into stimulation protocols that selectively enhance single-joint movements in neurorehabilitation.

## 1. Introduction

Spinal cord injury (SCI) is a life-altering event that leads to long-lasting motor impairment, and currently, there is no cure for paralysis (Armour et al., 2016; Center, 2021). Epidural spinal cord stimulation (SCS) has been gaining momentum as a neuromodulation intervention to restore movement to paralyzed areas below the injury (Angeli et al., 2018; Gill et al., 2018; Harkema et al., 2011; Minassian et al., 2007b; Rowald et al., 2022; Wagner et al., 2018). By selectively activating individual muscle groups at the appropriate phases of movement, epidural SCS can re-enable the performance of dexterous activities such as walking, cycling, swimming, and kayak paddling (Rowald et al., 2022; Wagner et al., 2018). However, the invasive nature of epidural SCS and the large group of experts needed with high associated costs may prevent epidural SCS from becoming an accessible therapy for the millions of people living with paralysis.

Transcutaneous SCS (tSCS) offers a promising non-invasive alternative to engage paralyzed muscles below the injury by activating the same neural structures via similar mechanisms as epidural SCS (Danner et al., 2011; Ladenbauer et al., 2010; Minassian et al., 2007a). Recent work suggests that it may be possible to achieve similar rehabilitative outcomes through this non-invasive approach (Al’joboori et al., 2020; Inanici et al., 2021; Samejima et al., 2022; Shapkova et al., 2020). However, the low selectivity of tSCS compared to epidural SCS (Hofstoetter et al., 2018) may limit the types and number of movements that can be enabled by it and, thus, its potential applications in exercise-based rehabilitation strategies. Moreover, the neural mechanisms behind functional enhancements in motor function enabled by tSCS remain poorly understood.

Electrode configurations that diverge from conventional tSCS have been shown to enable the preferential recruitment of either muscles ipsilateral to the stimulation site (Calvert et al., 2019) or rostral vs. caudal muscle groups (Krenn et al., 2015; Sayenko et al., 2015) in the lumbosacral region. Moreover, recent work has shown that multielectrode arrays have the potential to selectively target upper limb motor neuron pools in the cervical spinal cord (Calvert et al., 2019; de Freitas et al., 2022, 2021). In this work, we aimed to understand whether we could exploit the organization of motor neuron pools in the spinal cord to identify stimulation sites that result in optimal recruitment selectivity for different key leg muscles. We studied evoked compound muscle action potentials of lower-limb muscles to stimulation over the T10-L1 vertebral segments using conventional and multielectrode configurations of tSCS (**figure 1A,B**). First, we analyze recruitment curves for individual leg muscles (**figure 1C**) when the cathode is a single large electrode as in conventional tSCS (centered at T11/T12) or in one of six locations in a multielectrode configuration. We compare the recruitment selectivity enabled for each muscle at each cathode location and identify optimal electrode configurations that enable the highest selectivity for that muscle.

**Figure 1.**
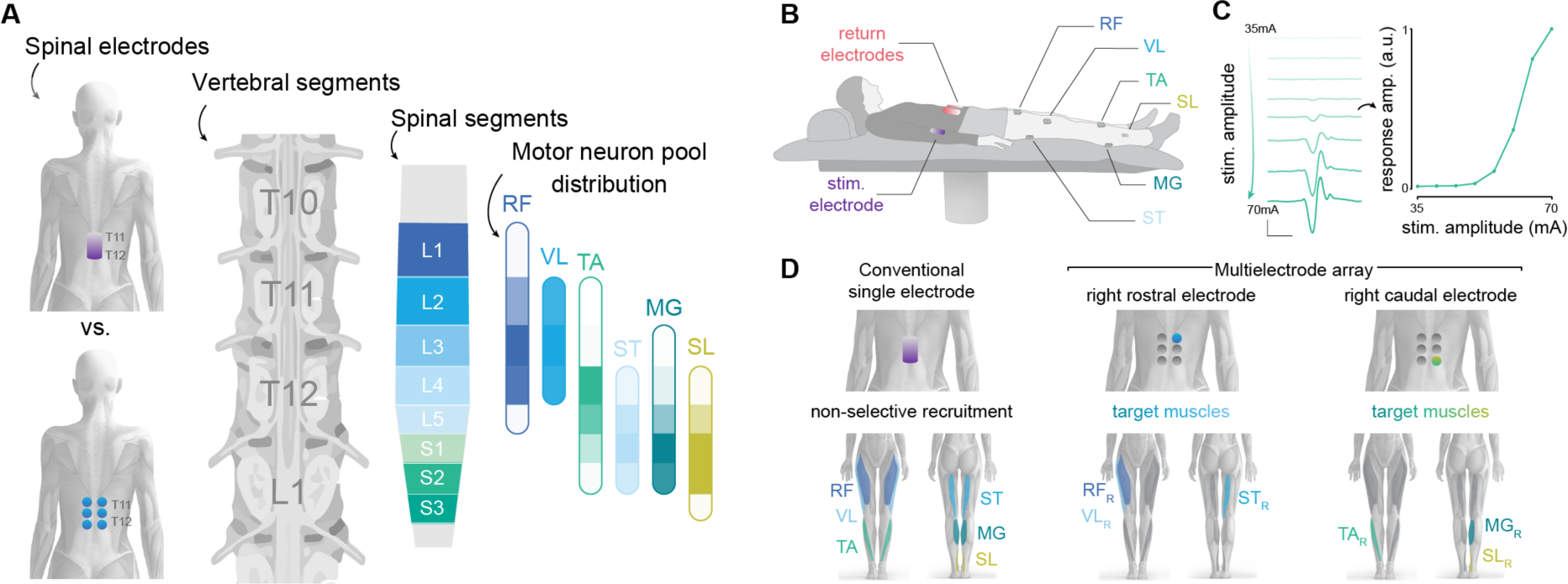
Experimental framework to compare muscle recruitment selectivity by conventional and multielectrode tSCS. (**A**) The rostrocaudal and unilateral organization of leg muscles’ motor neuron pools in the spinal cord may enable the selective recruitment of individual muscle groups by small diameter electrodes centered at the T11/T12 vertebrae. Modified from (Sharrard, 1964; Wagner et al., 2018). (**B**) We recorded leg muscle responses while pulses of tSCS were delivered at increasing stimulation amplitudes. (**C**) To quantify each muscle’s recruitment curve, we computed peak-to-peak response amplitude across a range of increasing stimulation amplitudes. Scale bars: 100 μV and 10 ms. (**D**) We hypothesize that compared to the non-specific recruitment by conventional single electrodes, a small diameter electrode positioned rostrally and over the right side would enable the selective targeting of right leg proximal muscles (hip, thigh), while a right caudal electrode would target right distal muscles (ankle). RF: rectus femoris; VL: vastus lateralis; ST: semitendinosus; TA: tibialis anterior; MG: medial gastrocnemius; SL: soleus; R: right leg.

Based on the rostrocaudal anatomical distribution of the motor neuron pools that innervate leg muscles (Hofstoetter et al., 2021b; Sharrard, 1964; Wagner et al., 2018) (**figure 1A**), we hypothesized that while conventional tSCS results in the non-selective recruitment of proximal and distal leg muscles, a right side cathode centered rostrocaudally at the T10/T11 interspinous ligament (overlapping the L1-L3 spinal segments) would primarily recruit proximal muscles in the right leg, whereas a right side cathode centered at the T12/L1 interspinous ligament (overlapping the L4-S3 spinal segments) would primarily recruit the right distal muscles (**figure 1D**).

Given that the probability of eliciting responses through direct recruitment of motor efferent fibers is dependent on the specific spinal segment and targeted muscle (Roy et al., 2012; Sayenko et al., 2015), it is essential to thoroughly investigate the activation mechanisms underlying potential enhancements in muscle recruitment selectivity. We hypothesized that the improved selectivity achieved through the multielectrode configuration would be mediated by the activation of muscle recruitment via sensory afferents, as evidenced by post-activation depression of the evoked response.

Finally, we combined the recruitment probability of a given muscle at a specific cathode site with previously reported segmental innervation probabilities at different spinal segments to validate our results and provide a neuroanatomical map of recruited spinal segments by each electrode configuration. Gaining a better understanding of the neural mechanisms behind these improvements in muscle recruitment selectivity by spatially selective tSCS may expedite the development of non-invasive technologies that can re-enable a broad repertoire of dexterous movements in rehabilitation and daily life.

## 2. Methods

### 2.1. Participants and experimental setup

We recruited 19 neurologically intact participants (9 female, 10 male, average age 23.12±4.34 years old, demographics in **supplementary table 1**) who gave their consent to take part in this study that was reviewed and approved by Washington University in St. Louis’ Institutional Review Board. Two participants (1 male and 1 female) withdrew from the study due to discomfort with the stimulation. One participant recruited early in the study was excluded from the analysis due to the use of monophasic, rather than biphasic stimulation, for a total of 16 participants included in the analysis for all figures.

### 2.2. Study Protocol

#### 2.2.1. Vertebral segment identification and electrode placement

We identified target vertebral segments T11/T12 via manual palpation, with validation by a second experienced research team member. The posterior iliac crest was first identified, and a transverse line was traced to the midline. The spinous process intersecting with this line was labeled with a surgical skin marker as the L4 spinous process. Interspinous ligaments were identified by palpation and labeled from L3/L4 to T9/T10. The skin around the midline from T10 to L2 was prepared with abrasive gel (NuPrep®, Weaver and Co. USA) using a Q-tips® in circular motions and wiped afterward with alcohol pads.

For the conventional tSCS condition, we positioned a single 5 x 9 cm rectangular electrode (all tSCS electrodes are PALS Neurostimulation Electrodes, Axelgaard Manufacturing Co., Ltd., USA) centered over the interspinous ligament of T11/T12 (**figure 1A**, top). Two interconnected 7.5 x 10 cm rectangular electrodes were placed on the abdomen bilaterally from the navel to serve as the return electrodes. All tSCS electrodes were treated with conductive spray (Signa® Spray, Parker Laboratories, Inc., USA).

Although palpation provides a useful starting point, anthropologic, clinical, and imaging studies have shown that variations in the number of vertebrae occur in 2-23% of the population (Bailey and Carter, 1938; Hahn et al., 1992; Akbar et al., 2010), and surface tSCS electrodes often need to be repositioned to achieve activation of the targeted muscles (Krenn et al., 2015). To ensure the accurate position of the conventional tSCS electrode over the lumbosacral enlargement, we conducted a verification process by assessing the balanced recruitment of proximal and distal muscles at comfortable stimulation amplitudes (refer to section *2.2.4 Recruitment curves*). If we observed large discrepancies in stimulation amplitudes required to recruit proximal vs. distal muscles, we adjusted the electrode placement by moving it one segment up or down as necessary. The horizontal midline of the adjusted conventional electrode placement was then used as the rostrocaudal center for the multielectrode configuration (middle electrodes).

In the multielectrode condition, we positioned six 3.2 cm diameter round electrodes 3 cm lateral to the midline and centered at the T11/T12 interspinous ligament (or center of conventional electrode if repositioned, **figure 1A**, bottom). Throughout the manuscript, we refer to the top, middle, and bottom row electrodes as the rostral, middle, and caudal electrodes, respectively. There was an approximate distance of 1-2 cm between electrodes in the longitudinal direction. In the multielectrode condition, only the abdominal electrode ipsilateral to the cathode was used as the return electrode.

#### 2.2.2. Data acquisition

Wireless electromyography (EMG) sensors (Trigno® Avanti, Delsys Inc., USA) were placed bilaterally according to SENIAM guidelines on the rectus femoris (RF), vastus lateralis (VL), semitendinosus (ST), tibialis anterior (TA), medial gastrocnemius (MG), and soleus (SL) muscles. The skin was prepared using the same procedure as with the spinal electrodes, and electrodes were repositioned if the baseline noise was larger than 10 µV. An additional wireless sensor (Trigno® Analog Input Adapter, Delsys Inc., USA) was connected via a BNC cable to the biphasic stimulator’s sync output for offline stimulation pulse alignment. Stimulation pulse triggering and amplitude were controlled via a data acquisition board (NI USB 6001, National Instruments, USA). EMG data was amplified using a data acquisition system (Trigno® Avanti Research+, Delsys Inc., USA; gain: 300; bandwidth 20-450 Hz), recorded at a sampling frequency of 2,000 Hz, and displayed in real-time using a custom-built software written by our group in Python (v3.10). EMG data were additionally bandpass-filtered offline between 10-1,000 Hz using a non-causal 2^nd^ order Butterworth filter.

#### 2.2.3. Transcutaneous spinal cord stimulation

Paired pulse tSCS was delivered using an isolated constant current stimulator (DS8R, Digitimer Ltd., UK) with a pair of charge-balanced biphasic pulses of 1 ms per phase duration and an inter-stimulus interval of 33.3 ms using a train generator (DG2A, Digitimer Ltd.). We delivered pulses of increasing stimulation amplitude to detect the motor threshold and saturation amplitude. We defined the motor threshold as the amplitude at which we observed a ≥ 20 µV peak-to-peak response amplitude within a latency of 10-30 ms in any muscle. The saturation amplitude was defined as the amplitude at which we no longer saw an increase in response amplitude of the first recruited muscles or the maximum amplitude tolerated by participants, whichever was lower.

#### 2.2.4 Recruitment curves

We performed recruitment curve recordings by increasing stimulation intensity from 5 mA below the motor threshold to the saturation amplitude, with 8 steps between amplitudes for a total of 10 stimulation amplitudes. Four repetitions of double-pulse stimulation were performed at each amplitude. Threshold detection and recruitment curve recordings were repeated for each of the seven electrode configurations (one in conventional tSCS, six in multielectrode tSCS). The testing order was randomized for the multielectrode configuration.

### 2.3. Data processing and analysis

Offline data analysis was performed in custom-built software written in Python and MATLAB® (The MathWorks Ltd., USA). Evoked responses were averaged across repetitions, with one average response waveform for each of the 10 stimulation amplitudes. Evoked responses for each muscle were normalized to the maximum averaged evoked response in both electrode configurations (one normalization for conventional tSCS and one normalization across the six round electrodes). Data was normalized separately for each condition since the maximum attainable response amplitude for each muscle was determined by either saturation or discomfort at that electrode configuration. To validate the robustness of our results to the normalization process, data was re-analyzed with one normalization across all conditions.

#### 2.3.1. Selectivity index

To quantify the degree of muscle recruitment selectivity enabled by each electrode position, we first generated a recruitment curve for each muscle. This curve represented the muscle’s peak-to-peak response as a function of stimulation amplitude. We then quantified the recruitment for each muscle as the area under the curve (AUC) of its recruitment curve normalized to the maximum possible recruitment i.e. maximal recruitment at all stimulation amplitudes. Last, we computed the selectivity *SI* of muscle *m* as the normalized recruitment *RECm* of that muscle minus the average normalized recruitment of *M* - 1 muscles, excluding the considered muscle, *m*. Conceptually, the selectivity index reflects the normalized recruitment of a given muscle compared to the average activation of all other muscles (Badi et al., 2021):

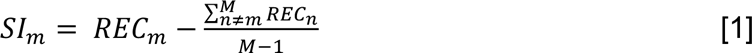

The selectivity index varies from-1 to 1 and reflects the activation of a targeted muscle compared to the average activation of all other muscles, with a value of 0 indicating that a given muscle is recruited equally to the average of all other muscles. This computation was performed for each muscle, electrode configuration, and participant.

To identify the target electrode for each muscle, we first compared the selectivity index enabled by three electrode positions ipsilateral to the target muscle (rostral: T10/T11, middle: T11/T12, caudal: T12/L1) and selected the electrode with the highest median selectivity index as the target electrode for that muscle. We then tested whether this target electrode enabled a significantly higher selectivity index for the targeted muscle compared to the conventional tCSS electrode and whether the selectivity index by the target electrode was higher than by its contralateral counterpart i.e. the small-diameter electrode at the same rostrocaudal level on the contralateral side.

#### 2.3.2. Paired pulse suppression

The amount of suppression by post-activation depression was calculated as the ratio between the second (R2) to the first (R1) response amplitudes. An R2/R1 ratio lower than 1 indicates that the amplitude of the evoked response to the second pulse was lower than the amplitude of the first response. The amount of suppression has been shown to be inversely related to the stimulation amplitude (Danner et al., 2016; Skiadopoulos et al., 2022), so we quantified suppression at the highest stimulation amplitude, where the amount of suppression should be the lowest.

#### 2.3.3. Recruitment probability and spinal activation maps

The probability of recruiting a given muscle was computed for each electrode configuration as the normalized AUC of its recruitment curve at that electrode position. AUC values for each muscle were normalized to the AUC in the electrode that achieved the highest level of recruitment. We then computed the recruitment probability for each muscle as the average normalized recruitment across participants. In this computation, a recruitment probability of 1 would indicate that that muscle was maximally recruited at that electrode position for all participants.

Recruitment probability estimates were used to model the activation of motoneuron pools in each spinal segment *Si* as a linear combination of the normalized leg muscle recruitment *Mj,* and *Wi,j*,, the expected segmental distributions of motoneuron pools innervating muscle *j* at spinal segment *Si,* (Wagner et al., 2018):

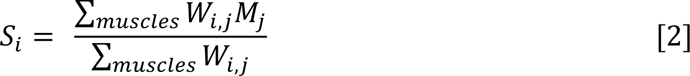

The coefficients *Wi,j* were obtained from an innervation probability map constructed by Hofstoetter et al., which included qualitative and quantitative data from thousands of participants in anatomical textbooks and electrophysiological studies (Hofstoetter et al., 2021b). Probability estimates for the VL muscle were derived from a separate anatomical study showing similar innervation in quadriceps muscles within the L2-L4 spinal segments (Sharrard, 1964; Wagner et al., 2018). The resulting spinal activation values were interpolated and superimposed onto a 2D image of the human lumbosacral spinal cord.

### 2.4. Statistics

Statistical analyses were performed using the SciPy statistics toolbox (v1.8.1) for Python. Data for left and right leg muscles were pooled together for analysis. Normal distribution was tested using the Kolmogorov-Smirnov test. A Friedman test for repeated measures was performed to determine the effect of the multielectrode rostrocaudal vertebral level (rostral, middle, and caudal) on selectivity index. A Wilcoxon signed-rank test with a Bonferroni correction for multiple comparisons was then used to determine significant differences in selectivity index across electrode positions. Separate Wilcoxon signed-rank tests were used to compare the selectivity index enabled by the target electrode against the conventional tSCS, as well as against its contralateral counterpart. A Wilcoxon rank sum test was used to test for significant suppression in each muscle by its target electrode.

## 3. Results

This study aimed to investigate the effects of multielectrode configurations of tSCS on muscle recruitment selectivity. We quantified evoked leg muscle responses to conventional and multielectrode tSCS and compared recruitment selectivity enabled by each electrode for a particular muscle. Our results show that rostrocaudal and ipsilateral positioning over muscle-specific motor neuron pools enhanced the recruitment selectivity of targeted muscle groups. The findings suggest that multielectrode tSCS has the potential to improve targeted muscle recruitment during functional movements, which could have important implications for motor rehabilitation in individuals with SCI.

### 3.1. Multielectrode configuration of tSCS results in enhanced rostrocaudal and unilateral selectivity compared to conventional tSCS

Overall, the motor threshold of the rostral and middle electrodes in the multielectrode configuration was higher than that of the conventional tSCS electrode, whereas the caudal electrodes had a lower motor threshold (**supplementary table 2**).

To study muscle recruitment enabled by conventional and multielectrode configuration electrodes, we recorded leg muscle responses over a broad range of stimulation amplitudes. Overlaid EMG responses to the first stimulation pulse at increasing amplitudes for the conventional and right caudal electrodes are shown for a representative participant in **figures 2A** and **B**. Notably, response latencies were progressively higher for caudally innervated muscles in both electrode configurations. This increase in latency is in agreement with previous observations (Courtine et al., 2007; Minassian et al., 2007a), suggesting that the increase in latency is due to the distance between motor neuron pools and the innervated muscle rather than the electrode location. Identification of the nature of the responses based on their latencies is not always definitive (Ertekin et al., 1996; Troni et al., 1996). Therefore, we applied existing neurophysiological methods to further analyze these responses.

**Figure 2.**
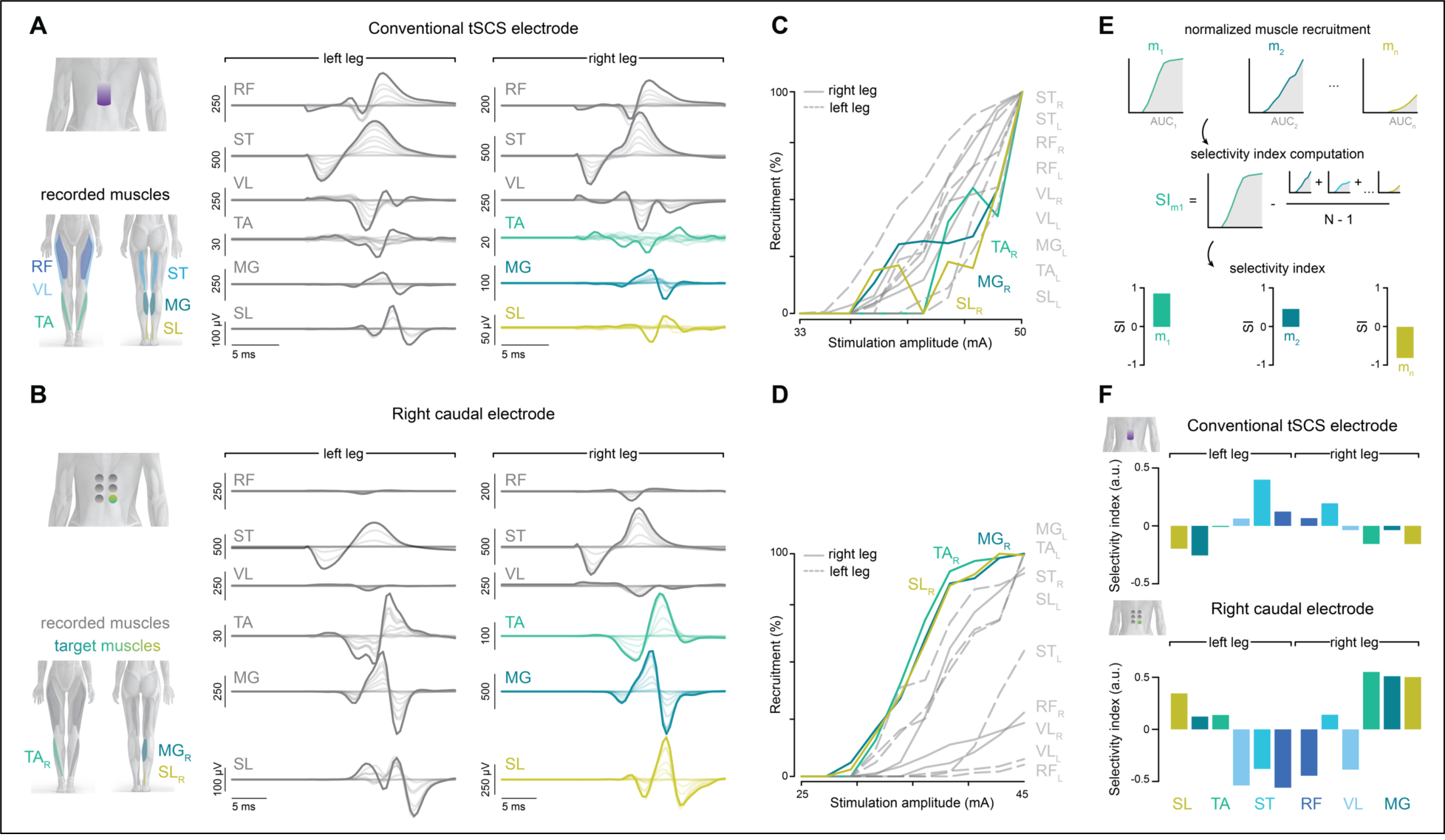
Muscle recruitment and selectivity by conventional and multielectrode tSCS. (**A**) Overlaid EMG responses are shown over a broad range of SCS amplitudes for the conventional electrode in a representative participant. Stimulation artifacts have been blanked for illustration purposes. (**B**) Overlaid EMG responses for the right caudal electrode in the same participant. (**C**) The EMG responses were averaged across *n* = 4 repetitions for each stimulation amplitude. The peak-to-peak amplitude was calculated to create a recruitment curve for each recorded leg muscle (color traces: targeted muscles). (**D**) Recruitment curves for muscle responses to stimulation by the right caudal electrode. (**E**) Illustration of selectivity index computation as the relative recruitment of a given muscle compared to all other muscles (for details, see *Methods*). (**F**) Selectivity index for all recorded muscles in conventional and right caudal electrode tSCS in this participant. RF: rectus femoris; VL: vastus lateralis; ST: semitendinosus; TA: tibialis anterior; MG: medial gastrocnemius; SL: soleus; R: right leg; L: left leg.

To understand the relationship between muscle recruitment and stimulation amplitude, we calculated the peak-to-peak amplitude of the averaged responses for each recorded leg muscle (**figure 2C,D**). The degree of muscle recruitment was proportional to the stimulation amplitude. However, the degree of recruitment for a given muscle at a given amplitude differed across electrode configurations. Stimulation by the conventional tSCS electrode configuration resulted in broad non-selective muscle recruitment, as reflected by similar motor thresholds across rostral, caudal, and bilateral muscles (**figure 2C**). In contrast, motor thresholds were lower in a subset of muscles recruited by the right caudal electrode, and their recruitment level was close to saturation by the motor threshold amplitude of the others (**figure 2D**, note the differences in recruitment level at ∼40 mA). The left leg distal muscles (TA, MG, SL) were also recruited at low amplitudes by the right caudal electrode in this participant. A common observation was that the targeted recruitment of a particular muscle group by the multielectrode configuration was usually accompanied by some undesired recruitment of the other unilateral muscle group (recruitment of right leg proximal muscles in the targeting of right leg distal muscles or *vice versa*), or the contralateral muscle group, as in this example. Therefore, we sought to quantify recruitment selectivity across muscles and electrode configurations.

To better interpret the degree of muscle recruitment selectivity enabled by each electrode position, we computed the selectivity index (Badi et al., 2021; Raspopovic et al., 2011) for all muscles (**figure 2E**). Selectivity indices for the conventional and right caudal electrodes in this participant are shown in **figure 2F**. The conventional tSCS electrode primarily enabled the recruitment of bilateral proximal muscles (RF, ST, VL), as indicated by a positive selectivity index. The area under the curve of the recruitment curve for distal muscles (TA, MG, SL) was lower than all others, resulting in a low and negative selectivity index for these muscles. In contrast, for the multielectrode configuration, the selectivity indices of distal muscles of the targeted (right) and non-targeted (left) legs were higher than most other muscles. Moreover, the relative activation of proximal muscles in both legs was lower than that for all other muscles, resulting in a low selectivity index for the non-targeted proximal muscles in this participant. To understand trends in recruitment selectivity that were common across participants, we performed group-level analyses.

### 3.2. Rostrocaudal and ipsilateral positioning over muscle-specific motor neuron pools enhances recruitment selectivity of targeted muscle groups

To identify the optimal electrode position to target each recorded leg muscle, we first computed the selectivity index for all muscles and electrode configurations across participants. We then compared the selectivity index enabled by each electrode rostrocaudal position (rostral, middle, or caudal) ipsilateral to the targeted muscle (**figure 3A**). Selectivity indices enabled by the multielectrode configuration depended on an electrode’s rostrocaudal position for most recorded muscles: Friedman effect of electrode’s rostrocaudal position in RF (*χ^2^ =* 9.75, *p* = 0.008), VL (*χ^2^ =* 15.06, *p* < 0.001), ST (*χ^2^ =* 0.063, *p* = 0.969), TA (*χ^2^ =* 10.94, *p* = 0.004), MG (*χ^2^ =* 6.06, *p* = 0.048), and SL (*χ^2^ =* 10.56, *p* = 0.005). Statistical significance for comparisons across electrode positions is shown above each muscle’s boxplot. The electrode with the highest median selectivity index was taken as the target electrode for that muscle for all subsequent analyses.

**Figure 3.**
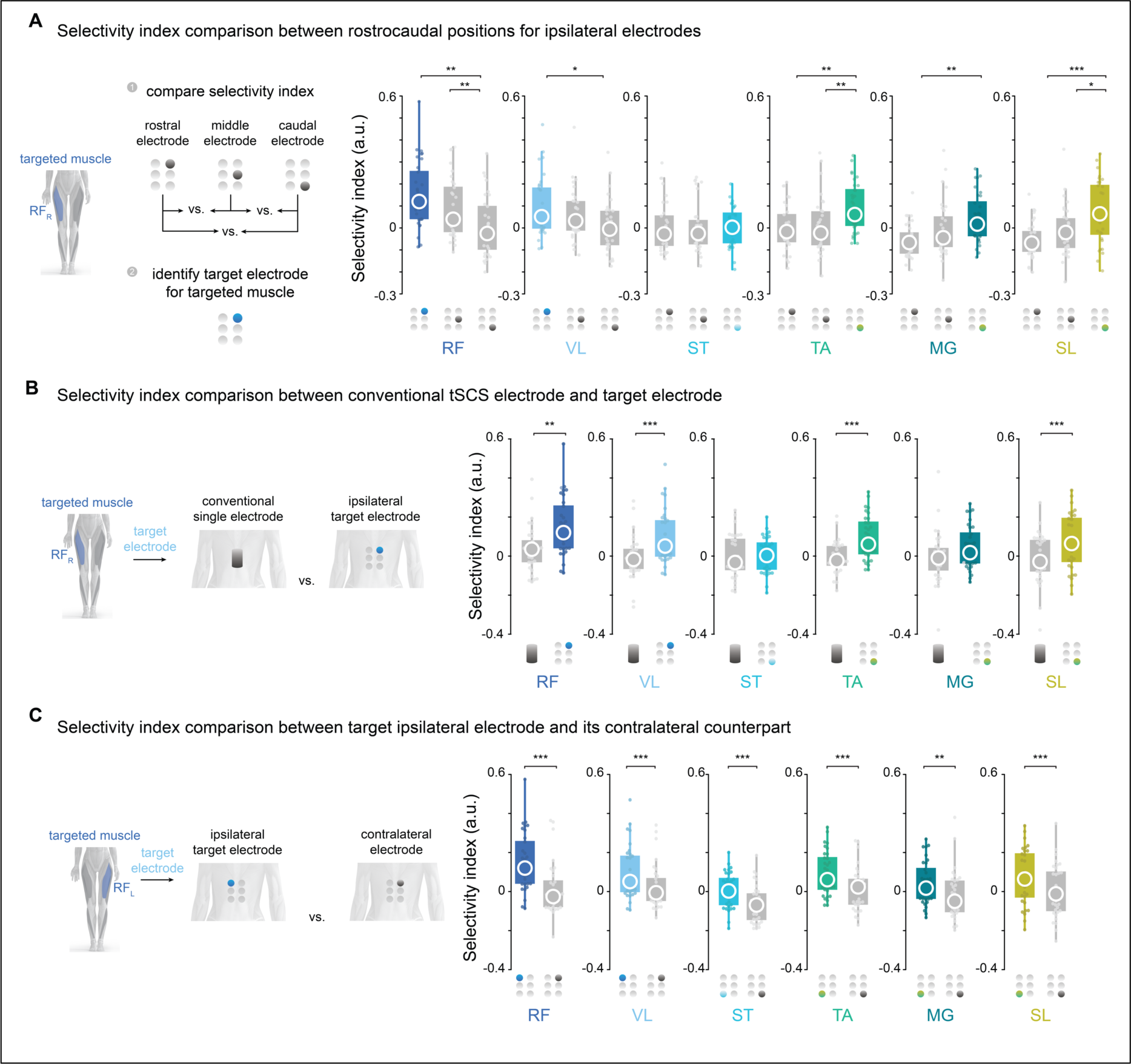
Selectivity index comparison between electrode types and positions. (**A**) Selectivity index comparison between electrodes in three rostrocaudal positions ipsilateral to the targeted muscle. The electrode with the highest median selectivity index is illustrated in color and was chosen as the target electrode for that muscle. Muscles from both legs are pooled together, with two data points per participant (e.g., one data point for the right RF targeted by the right rostral electrode and one data point for the left RF targeted by the left rostral electrode). (**B**) Selectivity index comparison between the target ipsilateral electrode and the conventional tSCS electrode. (**C**) Selectivity index comparison between the targeted ipsilateral electrode and its contralateral counterpart. Selectivity indices for most muscles were higher for electrodes ipsilateral to the targeted muscle than those achieved by the contralateral electrode. RF: rectus femoris; VL: vastus lateralis; ST: semitendinosus; TA: tibialis anterior; MG: medial gastrocnemius; SL: soleus; R: right leg; L: left leg. In the boxplots, the central circle is the median, while the top and bottom edges represent the 75^th^ to 25^th^ percentile, respectively. Friedman test for repeated measures for the effect of electrode position on selectivity index followed. Wilcoxon signed rank test with Bonferroni correction for multiple comparisons for comparisons across electrode positions. * p < 0.05; ** p ≤ 0.01; *** p ≤ 0.001.

We hypothesized that the target ipsilateral electrode in the multielectrode configuration could enhance the recruitment selectivity of a targeted muscle group compared to conventional tSCS (**figure 1D**). To test this hypothesis, we compared the selectivity index for each muscle enabled by the conventional tSCS and the ipsilateral target electrode (**figure 3B**). Overall, the target electrode in the multielectrode configuration significantly enhanced recruitment selectivity compared to the conventional electrode for most muscles: Wilcoxon signed rank effect of RF (*W =* 99, *p* = 0.002), VL (*W =* 43, *p* < 0.001), ST (*W =* 228, *p* = 0.500), TA (*W =* 54, *p* < 0.001), MG (*W =* 175, *p* = 0.096), and SL (*W =* 94, *p* = 0.001). Paired comparisons for individual muscles are shown in **supplementary figure 1**.

To test whether the improvements in muscle recruitment selectivity could be achieved by having the small diameter electrode at the right rostrocaudal level regardless of the electrode’s lateral position, we compared the selectivity index between the target ipsilateral electrode and its contralateral counterpart (**figure 3C**). As an example, we compared the selectivity index of the right RF when targeted by the right rostral electrode to the selectivity index of the same muscle targeted by the left rostral electrode. Overall, the ipsilateral target electrode significantly enhanced recruitment selectivity compared to its contralateral counterpart: Wilcoxon signed rank effect of RF (*W =* 54, *p* < 0.001), VL (*W =* 80, *p* < 0.001), ST (*W =* 55, *p* < 0.001), TA (*W =* 75, *p* < 0.001), MG (*W =* 100, *p* = 0.002), and SL (*W =* 65, *p* < 0.001).

Although selectivity index values for individual participants slightly varied when normalization of peak responses was performed across all electrode configurations (**supplementary figure 2**), the general trend across electrodes remained the same. The target electrode in the multielectrode configuration was the same for each muscle and achieved a higher level of selectivity than the conventional electrode and its contralateral counterpart. These results indicate that positioning small-diameter surface electrodes over the predicted locations of a muscle’s motoneuron pools in a rostrocaudal direction while maintaining laterality can improve the recruitment selectivity of that muscle.

### 3.3. Elicited responses with multielectrode configurations are mediated by posterior root-muscle reflexes

Studies examining the potential use of tSCS for recovery of motor function below the injury use the posterior root-muscle (PRM) reflex to confirm the effective position of the stimulating electrode over the spinal cord (Gad et al., 2017; Hofstoetter et al., 2015; Minassian et al., 2016; Sayenko et al., 2019). To investigate the reflex mechanisms behind spatially selective tSCS, we included in the recruitment curves protocol an additional paired pulse with a conditioning-test interval of 33.3 ms (**figure 4A**). We hypothesized that if elicited responses were indeed mediated by PRM reflexes, compound muscle action potentials to the test (second) pulse would reflect significant depression, a hallmark behavior of reflex responses (Hofstoetter et al., 2019; Olsen and Diamantopoulos, 1967).

**Figure 4.**
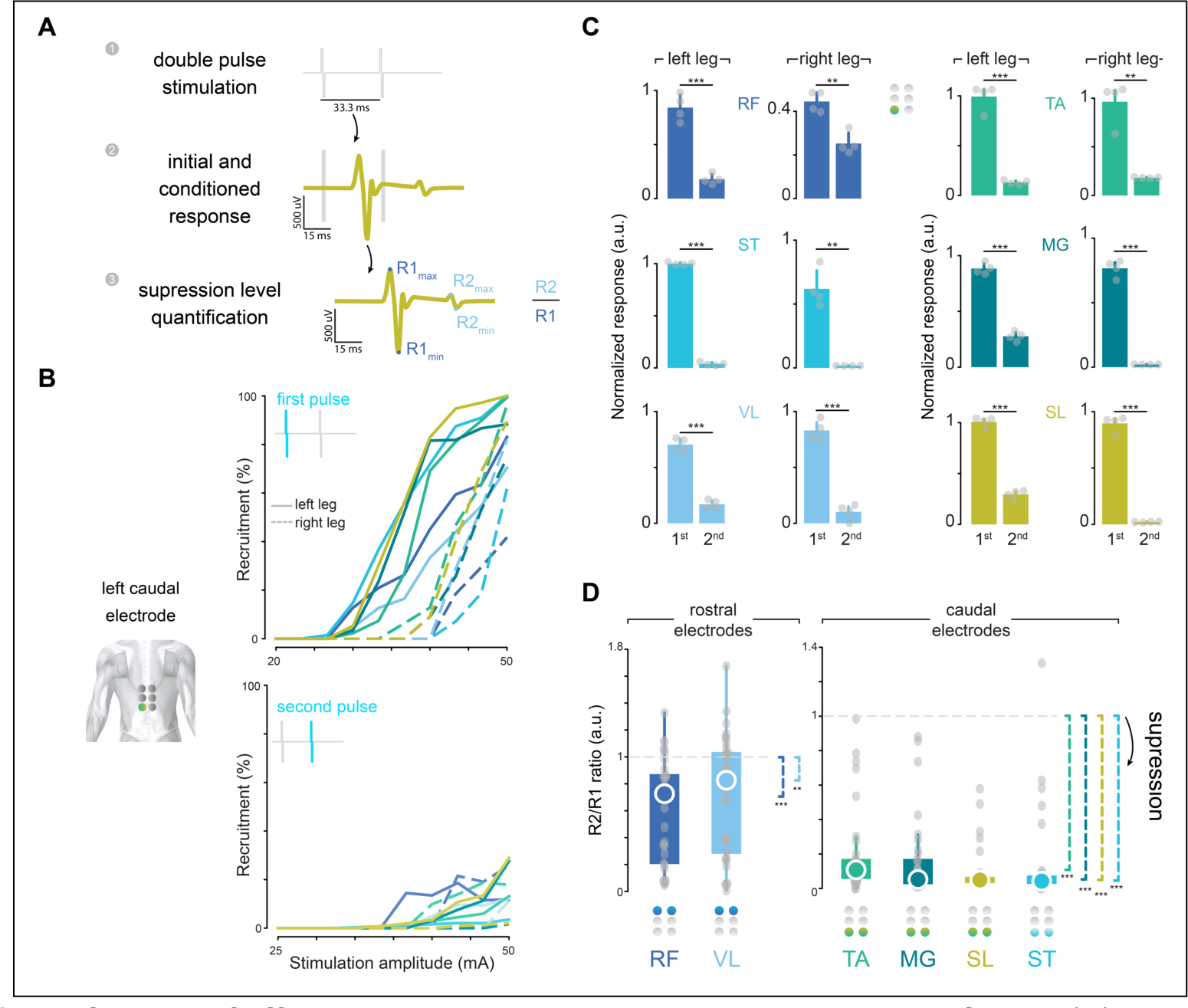
Verification of afferent stimulation by posterior root-muscle reflexes. (**A**) Exemplary paired-pulse response in SL muscle. We verified that small-diameter electrodes in the multielectrode configuration recruited primarily afferent fibers by the suppression of posterior-root muscle reflexes using a paired-pulse paradigm with an inter-pulse interval of 33.3 ms. The amount of suppression was quantified by the response amplitude ratio between the responses to the first and the second stimulation pulses. (**B**) Recruitment curves in a representative participant for the first and second pulse responses in the left caudal electrode from the multielectrode configuration. (**C**) First and second peak-to-peak responses at the highest stimulation amplitude in the same participant as in panel **B**. Peak responses are normalized to the highest muscle response across the multielectrode configuration. (**D**) Group analysis of R2 to R1 amplitude ratio across participants and electrodes. The right and left legs are pooled together. A value of 1 would indicate that the first and second responses had equal magnitudes, whereas a value < 1 would indicate that the response to the second stimulation pulse was lower than the one to the first pulse. The colored circle in the array indicates the optimal electrode for each muscle. In panel C, bars represent the mean ± s.d. Paired-sample *t*-test. In panel D, the central circle is the median, while the top and bottom edges represent the 75th to 25th percentile, respectively. One-sample Wilcoxon signed rank test with the alternative hypothesis that µ - 1 < 0. * p < 0.05; ** p ≤ 0.01; *** p ≤ 0.001. RF: rectus femoris; VL: vastus lateralis; ST: semitendinosus; TA: tibialis anterior; MG: medial gastrocnemius; SL: soleus; R: right leg; L: left leg.

Recruitment curves for the first (top panel) and second (bottom panel) pulses delivered through the left rostrocaudal electrode in a representative participant are shown in **figure 4B**. The amplitude of the response to the second pulse was largely suppressed in most muscles. However, the amount of suppression was inversely proportional to stimulation amplitude. Therefore, we quantified the level of suppression for each electrode at the highest stimulation amplitude, where the amount of suppression was generally the lowest. A comparison in response amplitudes between the first and the second pulse for the left caudal electrode in the same representative participant is shown in **figure 4C**. Overall, there was a significant reduction in response amplitude to the second stimulation by the left caudal electrode in that participant.

To analyze this effect across participants, we computed the level of suppression as the ratio between the first and second responses. A ratio smaller than one would indicate response suppression, i.e., that the response to the second stimulation pulse was lower than the response to the first pulse. The amount of suppression for each muscle, when targeted by its optimal electrode, is shown in **figure 4D**. There was a significant amount of post-activation depression of the evoked response by the paired-pulse paradigm at each optimal electrode configuration: one-sided 1-sample Wilcoxon signed rank test in RF (*W =* 489, *p* < 0.001), VL (*W =* 407, *p* = 0.004), ST (*W =* 527, *p* < 0.001), TA (*W =* 528, *p* < 0.001), MG (*W =* 528, *p* < 0.001), and SL (*W =* 528, *p* < 0.001). A significant amount of suppression was also observed in the conventional tSCS electrode (data not shown) and has been extensively reported by several groups (Dalrymple et al., 2023; Hofstoetter et al., 2021a, 2019; Oh et al., 2022). The amount of suppression for all multielectrode configurations and muscles is shown in **supplementary figure 3**. These results serve as a neurophysiological confirmation that the optimal multielectrode configuration enables the effective stimulation of the respective segmental afferents in the targeted leg muscles.

### 3.4. Recruitment probability for each muscle reflects the segmental distribution of its motor neuron pools in the spinal cord

To understand how different electrode configurations could target the posterior roots projecting to spinal cord segments containing the motor neurons responsible for activating the hip, knee, and ankle joints, we developed a neuroanatomical map of spinal cord activation for each electrode position. We first calculated the probability of activating each muscle at each electrode position by averaging the normalized recruitment of that muscle across participants (see *Methods*). Recruitment probabilities for electrodes located ipsilaterally and contralaterally to the targeted muscles at different rostrocaudal positions are shown in **figure 5A**. Next, we compiled an atlas of the expected anatomical locations for the motor neuron pools associated with the recorded leg muscles (Hofstoetter et al., 2021b; Sharrard, 1964; Wagner et al., 2018) (**figure 5B**). Finally, we projected the recruitment probability of each muscle onto its respective innervation probabilities at each spinal segment (**figure 5C**). The resulting spinal activation maps elicited by each multielectrode position are shown in **figure 5D**.

**Figure 5.**
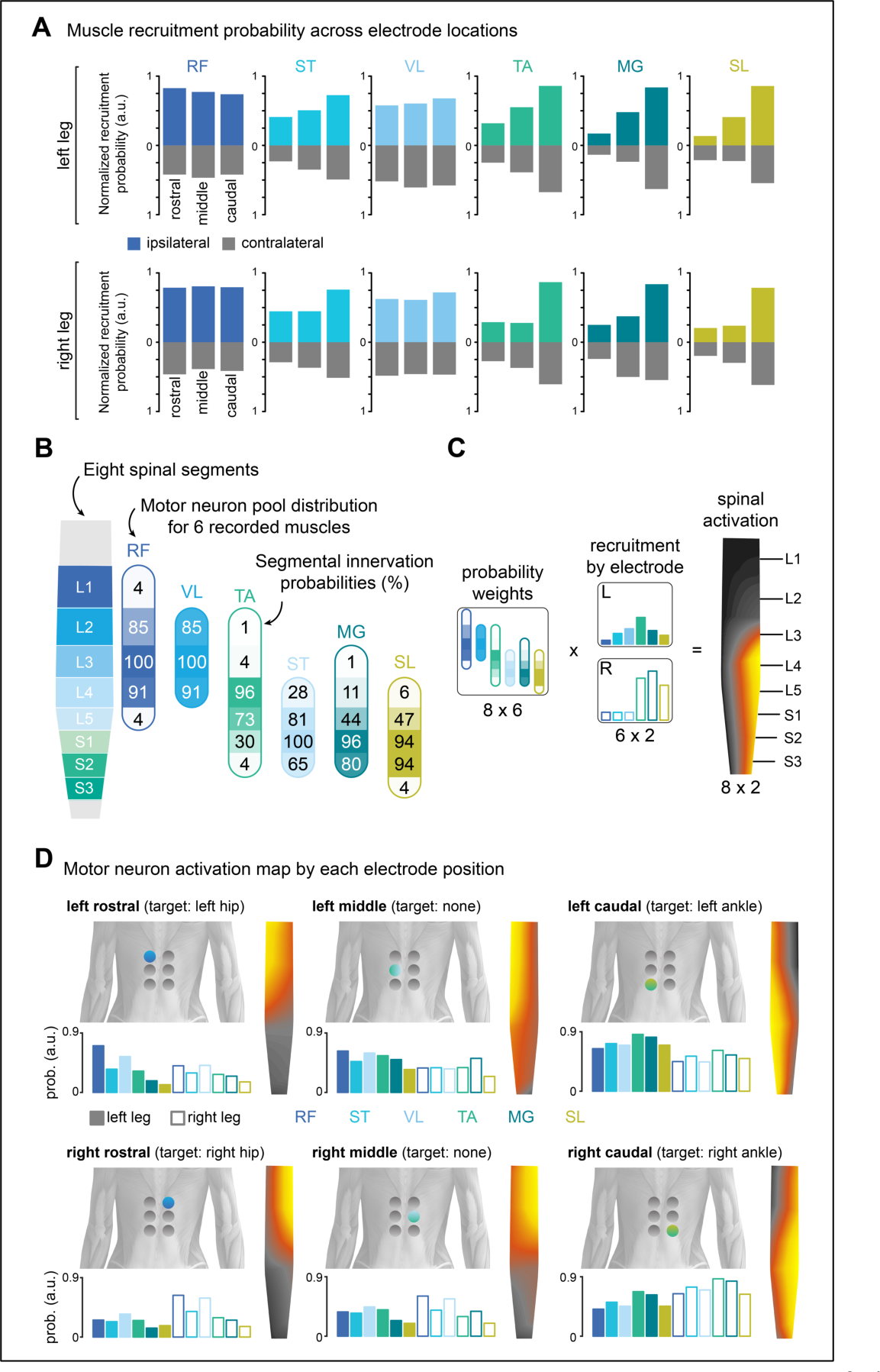
Neuroanatomical activation map by muscle recruitment probability. **(A**) Muscle recruitment probability for each muscle when the active electrode is at different rostrocaudal levels ipsilaterally (color) and contralaterally (grey) to the recorded muscle. Data were derived from the probability of recruiting a given muscle at a specific electrode position across all participants, where a probability of 1 would indicate that that muscle was maximally recruited for all participants at that electrode position. Note that values for recruitment probability by contralateral electrodes are also positive. (**B**) Segmental innervation probabilities for recorded leg muscles at different spinal segments. Innervation probabilities are reflected by percentage and opacity. Modified from (Hofstoetter et al., 2021b; Sharrard, 1964; Wagner et al., 2018). (**C**) Computation of spinal activation maps as a linear combination of the normalized recruitment of each muscle and its innervation probability at each segment. (**D**) Recruitment probabilities and motor neuron activation map enabled by each electrode configuration. RF: rectus femoris; VL: vastus lateralis; ST: semitendinosus; TA: tibialis anterior; MG: medial gastrocnemius; SL: soleus; R: right leg; L: left leg.

Our analysis of recruitment probability across participants revealed a high similarity between the optimal electrode configurations and the recruitment of the posterior roots projecting to the targeted spinal cord regions involved in the activation of the hip and ankle joints. For example, the left rostrocaudal electrode predominantly activated upper left lumbar segments (**figure 5D,** top left panel), while the right caudal electrode activated motor neuron pools located around right sacral segments (**figure 5D,** bottom right panel). It is important to note that the combination of the segmental innervation probabilities and the muscle recruitment probability was agnostic to the electrode configuration. Nevertheless, the spinal activation maps closely reflect the cathode location for that configuration.

## 4. Discussion

In this work, we present a technological framework to evaluate and quantify muscle recruitment selectivity enabled by different electrode configurations of tSCS. We identify optimal cathode positions to target different leg muscles by computing the selectivity index across 12 muscles and electrode configurations in 16 participants, with a total of 192 analyzed muscles. We demonstrate that a small-diameter multielectrode configuration of tSCS over the T10-L1 vertebral segments can enhance the rostrocaudal and unilateral recruitment selectivity of leg muscles compared to conventional tSCS using a single large electrode over the T11-T12 vertebra. Verification of post-activation depression suggests that these improvements in muscle recruitment selectivity are mediated by posterior root-muscle reflexes, which is a promising finding for the translation of this technology into clinical practice. Here, we discuss our findings and their implications in neurorehabilitation for people with neuromotor disorders.

### 4.1. Improving spatial selectivity by non-invasive technologies

Our study builds upon previous research in non-invasive tSCS, which has shown that stimulating different areas of the lumbar spinal cord can selectively activate motor neuron pools in leg muscles. Specifically, rostral and caudal positions of small electrodes centered over the midline have been shown to activate proximal and distal muscle groups, respectively (Krenn et al., 2015; Sayenko et al., 2015), and lateral positions can selectively recruit both proximal and distal muscles ipsilateral to the cathode position (Calvert et al., 2019).

In our work, we demonstrate that by combining these approaches, we can achieve selective activation of specific muscle groups in the hip, knee, and ankle joints, independently for the right and left legs. We found that different rostrocaudal positions of a small diameter electrode provided unique degrees of selectivity for different muscles. The rostral electrode, positioned lateral to the midline and centered rostrocaudally over the T10/T11 interspinous ligament, was the optimal position for RF and VL muscles, whereas the caudal electrode, centered over the T12/L1 ligament, was the optimal contact for TA, MG, and SL. However, optimal selectivity for the ST was more challenging to achieve. The caudal electrode was marginally better than other rostrocaudal positions, but this difference was non-significant. Despite the caudal segment being the most probable site for ST recruitment, we observed that ST muscle recruitment was highly likely to occur at low stimulation amplitudes across all three segments. This consistent recruitment of ST and other hamstring muscles at different stimulation sites has been previously observed by other groups and has been attributed to their broader segmental innervation than other muscles (Hofstoetter et al., 2021b). In their work, Hofstoetter and colleagues argue that PRM reflexes of the hamstring muscles may not provide useful information in intraoperative monitoring to guide electrode placement. Our results similarly suggest that responses from the other leg muscles should be prioritized when selecting the optimal electrode placement for a particular individual.

Interestingly, we found that the middle electrode, which was centered at the T11/T12 vertebral level, did not provide optimal selectivity for any muscle. This finding is noteworthy because conventional tSCS protocols often center the electrode at this level. We do not suggest that this placement is incorrect or ineffective, as if the objective were to target both proximal and distal muscles simultaneously by one stimulation site, T11/T12 or T12/L1 would be ideal placements. Rather, it may not be the optimal position for achieving selective activation of specific muscle groups.

Despite this, we do not recommend discarding the T11/T12 position in the multielectrode configuration altogether. In fact, it may be crucial in the context of multipolar configurations of tSCS, which have been shown to enhance recruitment selectivity further and prevent the unwanted activation of non-targeted muscles in epidural SCS (Struijk et al., 1993; Wagner et al., 2018; Hofstoetter et al., 2021b; Rowald et al., 2022). Therefore, while T11/T12 may not be the most selective electrode position for targeting specific muscles, it may still have important roles in optimizing tSCS protocols for further personalized improvements.

### 4.2. Toward personalized improvements in muscle recruitment

Our study revealed that achieving optimal recruitment selectivity using tSCS is not universal across individuals and targeted muscles. As shown in **supplementary figure 1**, the conventional tSCS electrode placement outperformed the target electrode in some participants’ muscles. While our results provide a valuable starting point for improving muscle recruitment selectivity in individuals with neuromotor disorders, it is important to recognize that interindividual differences in neuroanatomy and neurophysiology, particularly in the damaged nervous system, will require personalized approaches. To optimize stimulation protocols for each individual’s residual ability and specific needs, we believe it is crucial to test muscle recruitment for each participant and verify that these activations are mediated by posterior root-muscle reflexes. Modifications in stimulation parameters such as amplitude, frequency, and pulse width should then be made and reported based on these initial activation thresholds and selectivity starting points.

### 4.3. Importance of verification of effective stimulation sites by PRM reflex

The motor-enabling effects of SCS are primarily associated with the recruitment of proprioceptive afferent fibers in the posterior roots (Capogrosso et al., 2013; Dimitrijevic et al., 1980; Formento et al., 2018; Minassian et al., 2016). However, because of the complex topographic anatomy in the lumbosacral spinal cord (Wall et al., 1990), non-invasive stimulation at different lumbosacral vertebral segments can excite both afferent and efferent pathways to different degrees. The engagement of each pathway is crucial to both the immediate prosthetic effect that may be enabled by tSCS and the rehabilitation effect that may be observed through SCS-assisted therapy (Seáñez et al., 2022). Therefore, careful evaluation of evoked responses should be performed.

While direct recruitment of motor axons within the anterior roots, such as in functional electrical stimulation, can indeed produce desired movements in leg muscles Faghri et al., 1992; Popovic et al., 2011), the artificial recruitment of large-diameter motor fibers makes it technically challenging to produce and sustain large forces (Giat et al., 1993; Kirsch and Rymer, 1987). Moreover, the post-synaptic activation of motor fibers bypassing spinal and descending circuits limits the types of movements that can be performed to those that can be pre-programmed (Ajiboye et al., 2017), and prevents the voluntary modulation and neuroplasticity potential of these prosthetic effects. In contrast, the recruitment of afferent fibers by SCS enables interaction with descending and spinal circuits, which can enhance residual descending inputs (Minassian et al. 2016; Guiho et al. 2021) and lead to a natural recruitment order that is fatigue-resistant and capable of sustaining the whole body weight for extended periods (Formento et al., 2018; Wagner et al., 2018). This presynaptic recruitment of primary afferents, in turn, promotes neuroplasticity, which is believed to mediate the neurological recovery observed during neurorehabilitation facilitated by SCS (Asboth et al., 2018; Nishimura et al., 2013).

Overall, evoked responses by all target electrodes were mediated by PRM reflexes, as evidenced by the suppression of the conditioned response. However, the probability of efferent fiber recruitment differed substantially between L2-L4 innervated (RF, and VL) and L4-S2 innervated (ST, MG, TA, and SL) muscles (**figure 4D**). There were a few cases for proximal muscles in which the response average amplitude to the second pulse was higher than the average amplitude of the first response (values > 1 in RF, ST, and VL). This observation has been recently reported by others (Oh et al., 2022; Skiadopoulos et al., 2022), although, to our knowledge, has not been studied in detail. In contrast, there were no cases suggesting efferent recruitment in TA, MG, or SL, and only one case in ST. This suggests that higher stimulation amplitudes could be employed at caudal segments to target distal muscles with a reduced likelihood of recruiting motoneurons directly.

It should be noted that the degree of suppression was calculated at the maximum stimulation amplitude, which is associated with a greater likelihood of directly recruiting motor neurons. This selection was deliberate to facilitate a conservative analysis of recruitment mechanisms in a “worst-case” scenario. Routine applications of tSCS use stimulation amplitudes between 0.8 and 1.2 times the motor threshold to enhance motor function after paralysis (Hofstoetter et al., 2013; Inanici et al., 2021; Minassian et al., 2016; Samejima et al., 2022). As these amplitudes are considerably lower than saturation, we expect the probability of directly recruiting anterior roots in a therapy setting to be lower than those reported here. Nevertheless, because spine curvature can dramatically alter the probability of directly recruiting the anterior roots (Binder et al., 2021), verification of post-activation depression should be carefully confirmed in different therapeutic settings (e.g., sitting, standing, or in the position required to use a rehabilitation device).

### 4.4. Potential of improved selectivity in tSCS for neurorehabilitation

Despite the marked effects of stimulation frequency, intensity, and sensory feedback on evoked responses (Sayenko et al., 2015, 2014), continuous SCS is commonly used in studies involving SCS-assisted neurorehabilitation (Gerasimenko et al., 2015; Inanici et al., 2021; Meyer et al., 2020). In this application, stimulation parameters are typically fixed at the onset of therapy and kept constant across different types of movements. This continuous, non-selective SCS largely diverges from the natural spatiotemporal activation patterns observed during movement (Yakovenko et al., 2002) and can disrupt the natural feedback from proprioceptive afferents (Formento et al., 2018), which is critical to enable the spinal regulation of movements during fine motor control (Barra et al., 2022). And although long-term training combined with continuous SCS can indeed induce improvements in motor function that persist even without stimulation, they may take several months to a year of intense rehabilitation to appear (Angeli et al., 2018; Gill et al., 2018; Seáñez and Capogrosso, 2021).

Spatial and temporal control of SCS aims to facilitate movements through the selective activation of specific motor neuron pools at the appropriate phases of movement (Wenger et al., 2014, 2016). Recent clinical studies have shown that spatiotemporal epidural SCS can rapidly enable (within one week) powerful facilitation of walking, cycling, kayaking, and other dexterous activities in people with SCI (Rowald et al., 2022; Wagner et al., 2018) as well as upper limb movements and grasping ability in people with stroke (Powell et al., 2023).

Combining long-term activity-based training with tSCS has been shown to improve standing, balance, and induce functional recovery (Gad et al., 2017; Gerasimenko et al., 2015; Sayenko et al., 2019) that is comparable to that achieved through epidural SCS (Harkema et al., 2011; Rejc et al., 2015; Seáñez and Capogrosso, 2021). It is unlikely that non-invasive tSCS will achieve the same level of muscle recruitment selectivity as epidural SCS. However, using multielectrode configurations of tSCS to selectively target facilitation in certain muscles during movements could help individuals learn to perform complex movements faster, potentially accelerating their recovery process. By speeding up the recovery process, these technologies could then become more accessible to individuals with SCI by reducing the amount of training required and, ultimately, the cost.

### 4.5. Study limitations and implications for future studies

We show that muscle recruitment selectivity is higher in multielectrode configurations than in conventional tSCS when tested within the same session. However, in the context of rehabilitation, daily positioning of surface electrodes is necessary, making it impractical to test selectivity or PRM suppression on a daily basis. Therefore, it is important to investigate the consistency of our findings within and across participants when tested on different days. It remains unknown whether the optimal position for achieving selectivity in a particular muscle during one session will achieve the same level of selectivity when tested in a subsequent session. This is an important consideration for the development of effective and efficient tSCS-based rehabilitation protocols.

While our study demonstrated improved selectivity using single pulse responses, translating these findings into continuous muscle activation patterns required for movement poses significant challenges. During continuous stimulation, we may encounter difficulties in overcoming antidromic collisions between stimulation pulses and proprioceptive feedback, which can hinder movement ability (Formento et al., 2018). Stimulation tolerance is another important factor that may impact patient adherence and participant withdrawal (Moon et al., 2021). The higher charge density resulting from smaller electrodes may make it challenging to reach effective amplitudes near the motor threshold without causing discomfort. Further research is needed to investigate the neural mechanisms underlying muscle recruitment selectivity during continuous stimulation and to determine the optimal balance between efficacy and tolerability in tSCS.

Our study was conducted on neurologically intact individuals, while the intended population for this technology consists of individuals with SCI. An important consideration is that achieving selective stimulation in muscles with significant atrophy in people with SCI may pose a significant challenge. To enable selective stimulation in the atrophied muscle, we may require higher stimulation amplitudes that either cause discomfort or recruit non-targeted muscles before targeted ones. Therefore, it is essential to validate these findings in a population of individuals with SCI to ensure the feasibility of the technology in the indented population.

## 5. Conclusions

In summary, we demonstrate that the use of multielectrode tSCS results in enhanced rostrocaudal and unilateral selectivity compared to conventional tSCS. The optimal electrode position for targeting each leg muscle depends on the segmental innervation of its motor neuron pools within the spinal cord and allows for the targeted recruitment of specific muscle groups while minimizing the recruitment of other non-targeted muscle groups. As these are PRM-reflex-mediated responses, the interaction with descending and spinal neural circuits may enable the translation of this technology into effective therapies that aim to improve movement capacity in people with SCI.

## Data availability statement

Data from this study are available upon reasonable request from the authors.

## Author contributions

N.B. and J.D.P. real-time software platform development; N.B., I.S., and J.D.P., technological framework; N.B., R.H., J.F., L.L., R.K, and I.S. performed experiments; N.B., R.H., J.F., L.L., and R.K, and I.S. data curation, validation, and interpretation; N.B. and I.S. formal analysis; I.S. wrote original draft with N.B; N.B., R.H., J.F., L.L., and R.K, and I.S., reviewing and editing; I.S. study conceptualization, supervision, and securing funding.

## Supplementary Figures

**Supplementary Figure 1.**
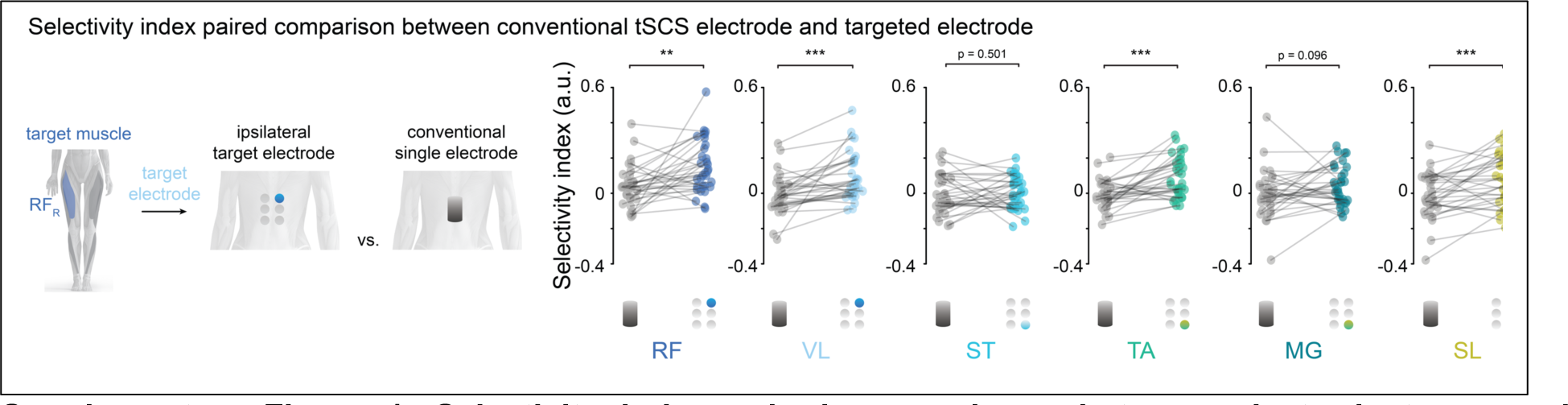
Selectivity index paired comparisons between electrode types and positions. Paired comparison in selectivity index between the targeted ipsilateral electrode and the conventional tSCS electrode. Same data as **figure 3B** with each muscle connected in both conditions. Contralateral muscles are pooled together for all panels. Wilcoxon signed rank test for comparisons across electrodes. * p < 0.05; ** p ≤ 0.01; *** p ≤ 0.001. RF: rectus femoris; VL: vastus lateralis; ST: semitendinosus; TA: tibialis anterior; MT: medial gastrocnemius; SL: soleus; R: right leg.

**Supplementary Figure 2.**
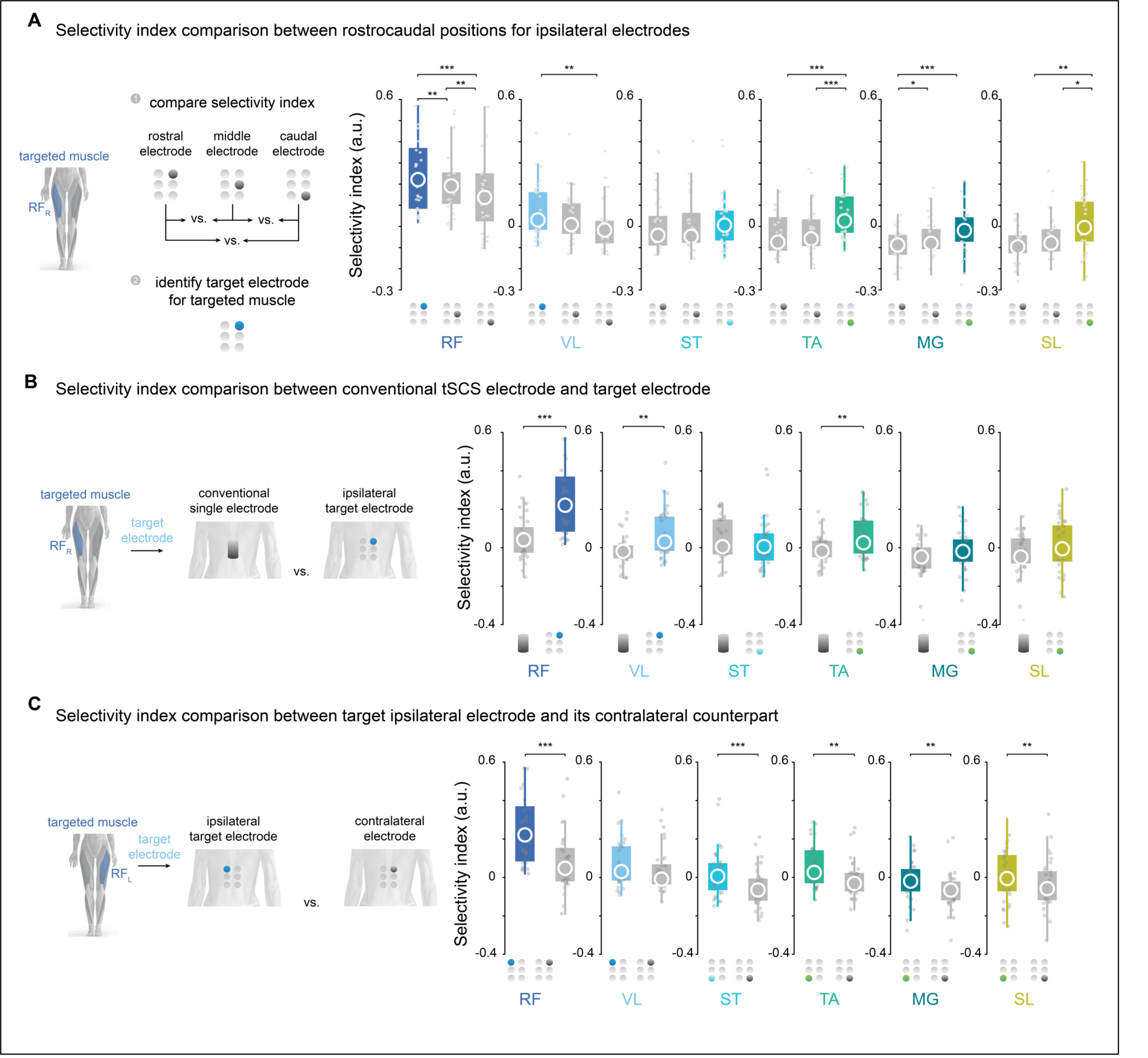
Selectivity index comparisons with normalization based on maximum absolute responses across all electrode configurations. Data processing and analysis for figure 3 was replicated, with the sole modification being that the normalization was performed across both electrode configurations rather than normalizing each muscle to the maximum response within each electrode configuration (one normalization for conventional tSCS and another normalization between the 6 contacts in the multielectrode configuration). (**A**) Selectivity index comparison between electrodes in three rostrocaudal positions ipsilateral to the targeted muscle. (**B**) Selectivity index comparison between the target ipsilateral electrode and the conventional tSCS electrode. (**C**) Selectivity index comparison between the targeted ipsilateral electrode and its contralateral counterpart. RF: rectus femoris; VL: vastus lateralis; ST: semitendinosus; TA: tibialis anterior; MG: medial gastrocnemius; SL: soleus; R: right leg; L: left leg. In the boxplots, the central circle is the median, while the top and bottom edges represent the 75^th^ to 25^th^ percentile, respectively. Outliers outside of scale are not shown for visualization purposes. Friedman test for repeated measures for the effect of electrode position on selectivity index followed. Wilcoxon signed rank test with Bonferroni correction for multiple comparisons for comparisons across electrode positions. * p < 0.05; ** p ≤ 0.01; *** p ≤ 0.001. RF: rectus femoris; VL: vastus lateralis; ST: semitendinosus; TA: tibialis anterior; MT: medial gastrocnemius; SL: soleus; R: right leg; L: left leg.

**Supplementary Figure 3.**
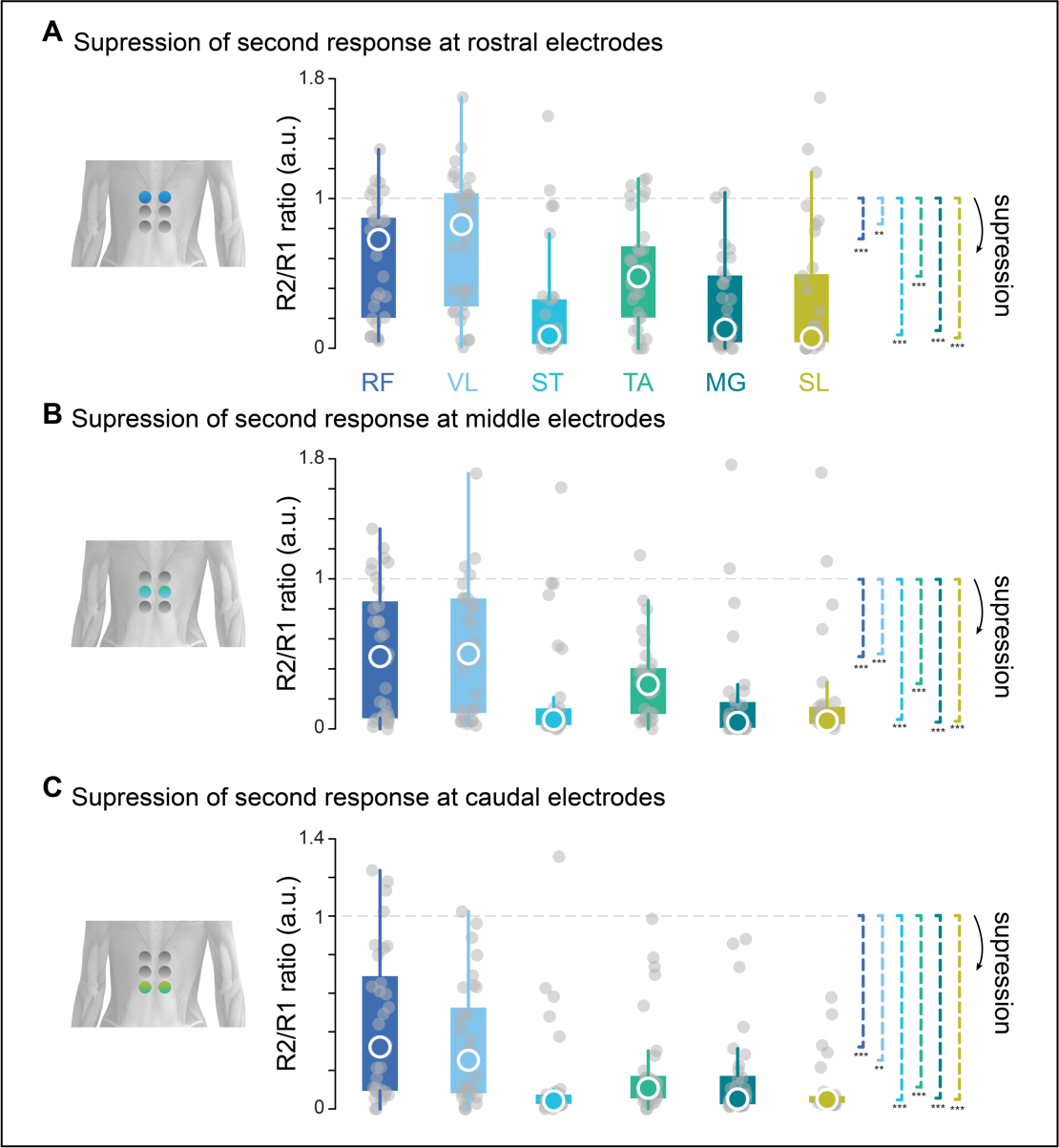
R2 to R1 amplitude ratio in all electrodes from the multielectrode configuration. (**A**) Group analysis of R2 to R1 amplitude ratio across participants and muscles for the rostral electrodes. The right and left legs are pooled together. (**B**) Suppression for middle electrodes. (**C**) Suppression for caudal electrodes. The central circle is the median, while the top and bottom edges represent the 75th to 25th percentile, respectively. One-sample Wilcoxon signed rank test with the alternative hypothesis that µ - 1 < 0. * p < 0.05; ** p ≤ 0.01; *** p ≤ 0.001. RF: rectus femoris; VL: vastus lateralis; ST: semitendinosus; TA: tibialis anterior; MG: medial gastrocnemius; SL: soleus; R: right leg; L: left leg.

**Supplementary Table 1.** Participant Demographics.

| Subj ID | Age (y) | Sex | Height (m) | Weight (kg) | Race | Ethnicity | Exclusion? |
| --- | --- | --- | --- | --- | --- | --- | --- |
| RC001 | 22 | Male | 1.80 | 99.79 | White | Non-Hispanic/Latino | N |
| RC002 | 23 | Male | 1.93 | 65.77 | White | Non-Hispanic/Latino | N |
| RC003 | 25 | Female | 1.75 | 104.30 | White | Non-Hispanic/Latino | N |
| RC004 | 22 | Male | 1.78 | 86.18 | White | Hispanic/ Latino | Withdrawn<br>(tolerability) |
| RC005 | 18 | Female | 1.55 | 51.3 | Asian | Non-Hispanic/Latino | N |
| RC006 | 22 | Male | 1.67 | 72.57 | White | Non-Hispanic/Latino | Monophasic<br>stimulation |
| RC007 | 25 | Male | 1.76 | 55 | Asian | Non-Hispanic/Latino | N |
| RC008 | 28 | Male | 1.65 | 131.54 | White | Non-Hispanic/Latino | N |
| RC009 | 22 | Female | 1.60 | 67.61 | White | Non-Hispanic/Latino | N |
| RC010 | 27 | Female | 1.52 | 97.52 | White | Hispanic/Latino | Withdrawn<br>(tolerability) |
| RC011 | 18 | Female | 1.52 | 61.23 | Asian | Non-Hispanic/Latino | N |
| RC012 | 25 | Female | 1.72 | 61.24 | White | Non-Hispanic/Latino | N |
| RC013 | 18 | Female | 1.57 | 52.16 | Asian | Non-Hispanic/Latino | N |
| RC014 | 20 | Female | 1.75 | 63.50 | White | Non-Hispanic/Latino | N |
| RC015 | 20 | Male | 1.93 | 99.79 | White/Asian | Hispanic/Latino | N |
| RC016 | 19 | Female | 1.55 | 65.77 | Asian | Non-Hispanic/Latino | N |
| RC017 | 24 | Male | 1.63 | 56.70 | Asian | Non-Hispanic/Latino | N |
| RC018 | 29 | Male | 1.85 | 78 | White | Non-Hispanic/Latino | N |
| RC019 | 34 | Male | 1.67 | 60 | White | Hispanic/Latino | N |

**Supplementary Table 2.** Average motor threshold for conventional and all rostrocaudal positions*.

| **Muscle** | **conventional**<br>**(mA)** | **left rostral**<br>**(x\*conv.)** | **right**<br>**rostral**<br>**(x\*conv.)** | **left middle**<br>**(x\*conv.)** | **right**<br>**middle**<br>**(x\*conv.)** | **left caudal**<br>**(x\*conv.)** | **right**<br>**caudal**<br>**(x\*conv.)** |
| --- | --- | --- | --- | --- | --- | --- | --- |
| L RF | 70.88 ± 23.68 | 1.11 ± 0.22 | 1.26 ± 0.17 | 0.99 ± 0.15 | 1.09 ± 0.14 | 0.89 ± 0.16 | 0.99 ± 0.21 |
| R RF | 69.64 ± 21.67 | 1.22 ± 0.2 | 1.09 ± 0.22 | 1.14 ± 0.14 | 0.98 ± 0.12 | 0.96 ± 0.18 | 0.87 ± 0.14 |
| L ST | 63.76 ± 21.23 | 1.34 ± 0.26 | 1.37 ± 0.16 | 1.12 ± 0.12 | 1.16 ± 0.16 | 0.92 ± 0.17 | 0.97 ± 0.2 |
| R ST | 65.0 ± 21.27 | 1.34 ± 0.19 | 1.26 ± 0.18 | 1.16 ± 0.19 | 1.12 ± 0.17 | 0.95 ± 0.19 | 0.89 ± 0.16 |
| L VL | 65.76 ± 22.78 | 1.12 ± 0.24 | 1.24 ± 0.20 | 0.98 ± 0.14 | 1.09 ± 0.15 | 0.88 ± 0.16 | 0.98 ± 0.24 |
| R VL | 69.55 ± 22.81 | 1.23 ± 0.24 | 1.11 ± 0.22 | 1.11 ± 0.15 | 1.00 ± 0.17 | 0.92 ± 0.19 | 0.88 ± 0.21 |
| L TA | 69.02 ± 22.67 | 1.27 ± 0.21 | 1.27 ± 0.19 | 1.07 ± 0.12 | 1.12 ± 0.12 | 0.85 ± 0.17 | 0.90 ± 0.2 |
| R TA | 68.67 ± 21.28 | 1.28 ± 0.15 | 1.25 ± 0.12 | 1.13 ± 0.15 | 1.13 ± 0.14 | 0.93 ± 0.21 | 0.87 ± 0.16 |
| L MG | 69.08 ± 26.09 | 1.32 ± 0.24 | 1.31 ± 0.20 | 1.07 ± 0.14 | 1.11 ± 0.17 | 0.86 ± 0.17 | 0.88 ± 0.21 |
| R MG | 63.9 ± 19.09 | 1.31 ± 0.18 | 1.27 ± 0.17 | 1.08 ± 0.11 | 1.06 ± 0.10 | 0.91 ± 0.20 | 0.88 ± 0.22 |
| L SL | 66.47 ± 25.91 | 1.37 ± 0.27 | 1.35 ± 0.19 | 1.11 ± 0.16 | 1.17 ± 0.14 | 0.88 ± 0.20 | 0.93 ± 0.21 |
| R SL | 66.24 ± 22.49 | 1.38 ± 0.25 | 1.29 ± 0.16 | 1.13 ± 0.17 | 1.09 ± 0.18 | 0.90 ± 0.19 | 0.87 ± 0.19 |

\*The average motor thresholds for all electrodes in the multielectrode configuration are shown as a ratio of the motor threshold with the conventional electrode.

